# Focal inbreeding and clonal transmission of *Plasmodium vivax* in pre-elimination Vietnam

**DOI:** 10.64898/2026.04.21.719798

**Authors:** Gabrielle C. Ngwana-Joseph, Nguyen Thi Huong Binh, Nguyen Thi Hong Ngoc, Jody Phelan, Susana Campino, Nguyen Quang Thieu, Taane G. Clark

## Abstract

Although malaria prevalence in Vietnam has markedly declined and the country presses closer to elimination, a highly focal reservoir of *Plasmodium vivax* malaria persists in its forested and border regions, threatening the country’s 2030 elimination targets. Genomic surveillance of malaria parasites can be a vital tool to produce fine-scale insights into the changing population dynamics of parasite populations, especially in population decline. Here, we analysed the genomic diversity of 18 newly sequenced *P. vivax* isolates from the Central Highlands, South Central Coast, and Southeastern regions of Vietnam collected in 2019, analysing them alongside 115 publicly available sequences from Vietnam collected between 2009-2016. Over these 10 years, infections became increasingly monoclonal, transmission remained highly focal, and the overall population structure was weak. Geographic factors, and not temporal factors, were a major driver of genetic substructure. Identity-by-descent (IBD) analyses revealed pockets of inbreeding in transmission hotspots, and high relatedness in parasites from within the same or adjacent provinces. Whilst within-population haplotype-based testing revealed minimal selection pressures on the 2009-2016 and 2019 populations, we observed multiple signals of differential selection of genetic variants involved in life-cycle specific processes. Overall, our work provides the most recent assessment of the genomic diversity of *P. vivax* in Vietnam, revealing relics of evolution in a parasite population in decline. Continued genomic surveillance, especially in outbreak contexts, and with more recent samples will be a crucial strategy to inform malaria elimination activities.

**Author Summary:** *Plasmodium vivax* is the most geographically widespread malaria parasite and the predominant species in the Greater Mekong Subregion. In Vietnam, *P. vivax* malaria continues to persist in remote forested settings and along its international borders. Using genomic approaches, we analysed 133 *P. vivax* genomes from the Central Highlands, South Central Coast, and Southeastern regions of Vietnam, including 18 newly sequenced isolates collected in 2019 and 115 publicly available genomes sampled between 2009 and 2016. Our work describes the evolutionary and genetic consequences of intensive control efforts that have led to decreasing transmission and population decline. We found decreasing infection complexity and increasing clonal inbreeding, which suggests localised transmission of clonal lineages in transmission hotspots. Over the 10-year sampling period, we observed consistent allele frequencies in putative drug resistance loci, suggesting an absence of drug-driven directional selection. Our findings highlight the importance of genomic surveillance in monitoring parasite persistence, which will be critical for guiding public health decisions to sustain progress towards malaria elimination.

## Introduction

Vietnam, along with countries in and around the Greater Mekong Subregion (GMS), is actively pursuing elimination of *Plasmodium vivax (P. vivax)*, the leading cause of malaria in the country, by 2030[1]. Between 2013 and 2023, *P. vivax* malaria cases in the country decreased by 97%, from 6,901 to 195, and there have been zero malaria deaths since 2019, representing the substantial success Vietnam has made in reducing its *vivax* malaria burden[2]. However, elimination of *P. vivax* malaria is complicated by the inherent nature of its biology, where some liver-stage parasites commit to a dormant, hypnozoite stage, which can cause disease relapse after primary infection and treatment[3]. Moreover, *P. vivax* infections are characterised by low parasitaemia[4], leading to a high occurrence of sub-patent and asymptomatic infections in countries with active transmission of *P. vivax*.

In Vietnam, the epidemiology of *P. vivax* malaria is complex, and residual active foci of transmission persist, threatening the elimination goals set by the World Health Organization’s (WHO’s) Mekong Malaria Elimination (MME) programme. Transmission of *vivax* malaria in Vietnam is highly focal, with intense transmission in the forested, mountainous, and hilly regions in the central and southern provinces. Over the past 5 years, reported cases are concentrated in 20 provinces, with the highest transmission in Gia Lai, Khanh Hoa, Lai Chau, Phu Yen, Binh Phuoc and Dak Lak. Moreover, malaria progressively and disproportionately affects poor ethnic minorities in remote areas[5,6], forest workers in labour-intensive industries (such as hydropower construction sites and plantations)[6], and forest-going mobile and migrant populations[7]. This presents a unique challenge in ensuring adequate coverage of effective interventions to the most malaria-vulnerable populations.

In near-elimination settings, proactive surveillance, rapid detection, and treatment of malaria cases are vital to suppress local transmission and prevent its re-establishment in regions of high receptivity. In this context, population genomics has been a powerful tool that has provided high-resolution insights into spatiotemporal trends in *vivax* transmission dynamics in local parasite populations across the world. These observations have been at micro-scales, within provinces[8,9] or countries[10], to macro-scales, across subregions[11–13] and the entire globe[14,15]. However, very few studies investigate the population genetics of *P. vivax* specifically in Vietnam. As *P. vivax* is sympatric with *P. falciparum* in the country, elimination efforts and population studies have been focused on *P. falciparum* due to the urgency posed by multidrug resistance, and the greater morbidity and mortality associated with *falciparum* malaria[16–18]. Studies that do investigate the population genetics of *P. vivax* in Vietnam generally use panels of microsatellite markers[19] instead of whole-genome sequencing (WGS) data. Using this approach is not representative of the full genomic landscape and biases the study of the genome to known hotspots, leaving the rest of the genome relatively unexplored. While WGS data is currently available for Vietnamese *P. vivax*[14][20], WGS analysis specific to the country has not been performed, and thus, our understanding of *P. vivax* population dynamics in Vietnam is limited.

Vietnam is therefore a unique setting to assess the changing transmission dynamics and selective pressures on the parasite population using WGS, especially as the country makes the critical transition towards malaria elimination. Here, we performed WGS of *P. vivax* cases from three high-transmission provinces in Vietnam. We characterised infection clonality, population structure, ancestry, relatedness, and identified loci of selection pressure on the parasite’s genome. Finally, we performed comparative analysis of the newly sequenced genomes with publicly available sequences[20] to make inferences about spatiotemporal trends in evolution and transmission of *P. vivax* to help guide interventions targeting this sophisticated and highly resilient parasite.

## Methods

### Study Sites and Sample Collection

Parasite DNA was harvested in the form of dried blood spots for study from 44 microscopy-confirmed symptomatic *P. vivax*-infected patients, conducted by trained field researchers in Vietnam based at the National Institute of Malariology, Parasitology, and Entomology (NIMPE) in Hanoi, Vietnam, in 2019. Samples were collected from the Southeast region, represented here by Ho Chi Minh and Binh Phuoc, the South Central Coast region, represented by Khanh Hoa, and the Central Highland region, represented by Gia Lai.

### Ethics Statement

All patients provided written or verbal informed consent, approved by the research ethics committee of NIMPE, Ministry of Health, Vietnam (Ref. 1096/QD-VSR).

### Whole-genome Sequencing

DNA was extracted from dried blood spots using a DNeasy Blood and Tissue Kit (Qiagen) and amplified using an established selective Whole Genome Amplification (sWGA) protocol to preferentially amplify *P. vivax* DNA prior to sequencing[21]. Briefly, for each sample, a unique sWGA reaction was performed using 30 μL of extracted DNA, corresponding to between 5 ng and 10 ng of DNA. This was combined with 5 μL of Phi29 DNA Polymerase Reaction Buffer (New England BioLabs), 0.5 μL of Bovine Serum Albumin (New England Biolabs), 0.5 μL of SWGA primers (2.5 μM)[21], 5 μL dNTPs (New England Biolabs), and water to a total reaction volume of 50 μL. The SWGA reactions were then carried out in a thermocycler using the following programming steps: 5 minutes at 35°C, 10 minutes at 34°C, 15 minutes at 33°C, 20 minutes at 32°C, 25 minutes at 31°C, 16 hours at 30°C, 10 minutes at 65°C to degrade the Phi29 enzyme, and a final hold at 4°C. Isolates were then prepared for sequencing using the KAPA Pure Beads (Roche) Genomic DNA purification protocol. Samples containing a minimum of 300 ng DNA WGS (27/44 (61.4%)) were sequenced on an Illumina Novaseq 6000 platform at Eurofins Genomics, Germany, producing a minimum of 5M paired reads (150 bp reads) per sample.

### Sequence Processing, Variant Calling, and Quality Control

All raw Illumina sequence reads for the successfully sequenced 27 Vietnamese isolates, along with the 724 publicly available isolates[20] from Southeast Asia retrieved from the European Nucleotide Archive (ENA) in fastq format, were processed using established pipelines[15]. Raw reads were trimmed using *trimmomatic* (v0.40)[22] with parameters PE -phred33, LEADING:3, TRAILING:3, SLIDINGWINDOW:4:20 MINLEN:36 to remove low quality sequences. Trimmed reads were aligned to the PvP01_v1 reference genome[23] using *bwa-mem* (v.0.7.17, default parameters)[24], producing a BAM file for each sample. The *samtools* functions fixmate and markdup (v1.22, default parameters)[25] were applied to the resultant BAM files. *GATK* (v4.1.4.1)[26] functions BaseRecalibrator and ApplyBQSR functions were used to improve BAM files for all isolates, using a previously published training set of high-quality *P. vivax* SNPs to calibrate variant calling. *GATK* (v4.1.4.1) function HaplotypeCaller (parameters: -ERC GVCF) was used to call SNPs and indels, producing individual sample VCFs with validated variants. Validated VCFs were imported into GenomicsDB using the *GATK* function GenomicsDBImport (v4.1.4.1). The *GATK* function GenotypeVCFs (v4.1.4.1) was used to produce a combined, multi-sample VCF: one containing SNP variants only and one containing indel variants only. The resulting VCF files contain genotype calls for all variant sites across all samples. Only variants in the core, non-hypervariable regions of the *P. vivax* genome were included and assigned a Variant Quality Score Log-Odds (VQSLOD) score using *GATK* functions Variant Quality Score Recalibration and ApplyVQSR (v4.1.4.1). Variants with a VQSLOD >0, meaning they are more likely to be false variants than true variants, were excluded. Looking at SNPs exclusively, isolates with >40% of SNPs with missing genotype calls were excluded from further analysis. Mixed call SNPs were designated genotypes determined by a ratio of coverage in which nucleotide calls were 80% or higher. Finally, monomorphic and multiallelic SNPs were excluded, leaving a final dataset comprising 514,951 high-quality, bi-allelic SNPs across 615 samples from Vietnam (N=133), Cambodia (248), China (16), Laos (2), Myanmar (28), North Korea (6), and Thailand (182). Downstream coding effects of SNPs were predicted using SnpEFF[27].

### Population structure, ancestry, and infection clonality

Sample geography was visualised in *R* (v4.4.3) using the *rnaturalearth* (v1.1.0), *rnaturalearthdata* (v1.0.0), *sf* (v1.0_20) and *ggplot2* (v3.5.2) packages. Population structure of isolates was inferred from a binary matrix of pairwise Manhattan distances between isolates generated using PLINK (v1.90b6.21)[28] and visualised in R (v4.4.3) using *ggplot2* (v3.5.2). This binary matrix was also used to construct a neighbour-joining tree using the R package *ape* (v5.8_1) and visualised in iTOL[29]. Infection clonality was determined by calculating F_WS_, a measure of within-host diversity, using moimix with an allele frequency threshold of 0.1% (https://github.com/bahlolab/moimix). *ADMIXTURE* software (v1.3.0)[30] was used to estimate ancestry in unique individuals. Analysis was conducted on a filtered set of 126,019 SNPs, with minor allele frequencies (MAFs) of at least 1%. *PLINK* (vv1.90b6.21) was used to convert this subset of SNPs from a VCF to a BED file. The most probable number of ancestral populations, K, was estimated through cross-validation error of 1-10 dimensions of eigenvalue decay and visualised in *R* (v4.4.3) using *ggplot2* (v3.5.2).

### Identity by descent and detecting recent positive selection

Pairwise relatedness of isolates was estimated through identity by descent (IBD) analysis using *hmmIBD* (https://github.com/glipsnort/hmmIBD)[31]. *hmmIBD* employs a hidden Markov model to haploid genotypes to identify tracts of extended haplotype homozygosity. IBD analyses were exclusively performed on monoclonal isolates (F_WS_ ≥ 0.95) in country-level populations ≥10, and on biallelic SNPs with MAF ≥1%. The fraction of genomic sites sharing IBD between isolates was plotted as a network in R (v4.4.3) using *igraph* (v.0.10.16). Nodes represent unique isolates and edges between two nodes represents the fractional IBD sharing between any two isolates based on relatedness thresholds equivalent to clones (IBD ≥ 0.95), full siblings (IBD ≥ 0.475), half siblings (IBD ≥ 0.2375) and quarter siblings (IBD ≥ 0.1188), as well as overlaid on a map of Vietnam and visualised using R’s *ggplot2* (v3.5.2).

Signals of recent positive selection were detected using the R package *rehh* (v3.2.2)[32] on monoclonal isolates (F_WS_ ≥ 0.95) and on biallelic SNPs with MAF ≥1%. The *rehh* package calculates the integrated haplotype score (*iHS*) and the standardised log ratio of integrated haplotype homozygosity (*Rsb*). The *iHS* statistic detects selection within a population, while *Rsb* identifies selection between populations. Using 10 kb sliding windows with at least 5 SNPs, critical loci were defined as having an *iHS* value > 4.0 (− log_10_[1 – 2 | Φ_iHS_ – 0.5 |] > 4.0) and (-log_10_[p-value]) > 5.0 for *Rsb*. These cutoffs were determined using a Gaussian approximation method. Finally, to identify SNPs that drive allele frequency differences between subpopulations, python (v3.13.5)’s scikit-allel (v1.3.13) was used to calculate Hudson’s F_ST_. Hudson’s estimator was preferred here as it is less biased by unequal sample sizes between populations compared to Weir and Cockerham’s estimator.

## Results

### Sample selection and genome data

We collected and sequenced 27 *P. vivax* clinical isolates in 2019 across the Central Highlands, South Central Coast, and Southeast regions of Vietnam. Of these, 18 successfully passed data quality-control and were retained for population genetic analysis. These 18 isolates were sourced from Vietnam’s Gia Lai (11) and Khanh Hoa (7) provinces, with a median genome-wide coverage of 48.03-fold (**Table S1**). After processing with publicly available sequence data from Vietnam (133) and countries in and around the GMS (Cambodia (248), China (16), Laos (2), Myanmar (28), North Korea (6), Thailand (182) (**Table S2**), we obtained a dataset of 615 samples encompassing 514,951 high-quality biallelic SNPs in the core, non-hypervariable regions of the *P. vivax* genome. Of these SNPs, 249,392 (48.4%) were in coding regions, and 96,296 (18.7%) were missense (non-synonymous) variants. Vietnamese isolates were grouped by region for spatial analysis: Southeast (N = 83), represented here by Ho Chi Minh and Binh Phuoc, the South Central Coast (N = 8), represented by Khánh Hòa, and the Central Highlands (N = 38), represented by Gia Lai.

### Genomic complexity of P. vivax infections in Vietnam has reduced over time

Within-host diversity of *P. vivax* isolates from Vietnam was estimated using the fixation index, where an F_WS_ ≥0.95 is indicative of an infection predominated by a single parasite clone. High transmission intensity is associated with higher infection multiplicity (F_WS_ <0.95). Initially, the population appeared to be multiclonal in nature, with a mean F_WS_ of 0.88 and only 69.6% (80/133) of isolates with an F_WS_ ≥0.95. However, stratifying samples by year of sample collection revealed that over time, F_WS_ has been trending upwards, indicative of decreasing infection complexity (**Fig. S1**). In the 2009-10 population (N = 14), mean F_WS_ was 0.89, slightly decreasing to 0.86 in 2014-15 (N = 67), before sharply increasing to 0.99 in 2019 (N = 18). Despite the increase in F_WS_ between 2009 and 2019, the difference was not statistically significant (Wilcoxon rank-sum test, W = 90, p = 0.174).

### Vivax from the GMS forms two major population clusters

The genetic relationship between *P. vivax* parasites in and around the GMS (**Fig. 1a**) was initially explored through a SNP-based principal component analysis (PCA) (**Fig. 1b**) and accompanying neighbour-joining tree (**Fig. 1c**), revealing that population substructure was strongly influenced by geographic provenance. The newly sequenced isolates from Vietnam’s Gia Lai (11) and Khanh Hoa (7) provinces clustered with all publicly available Vietnamese sequences, showing close genetic similarity between circulating parasite populations within the country. Vietnamese, Cambodian, and Laotian populations clustered together, forming a distinct population centre, whereas Thai, Burmese, Chinese and North Korean isolates formed a secondary population cluster in a looser grouping, exhibiting greater structural diversification and genetic divergence. These findings were recapitulated in an accompanying neighbour-joining tree, where Cambodian, Vietnamese, and Laotian isolates separated distinctly from all other populations (**Fig. 1c**).

**Figure 1.**
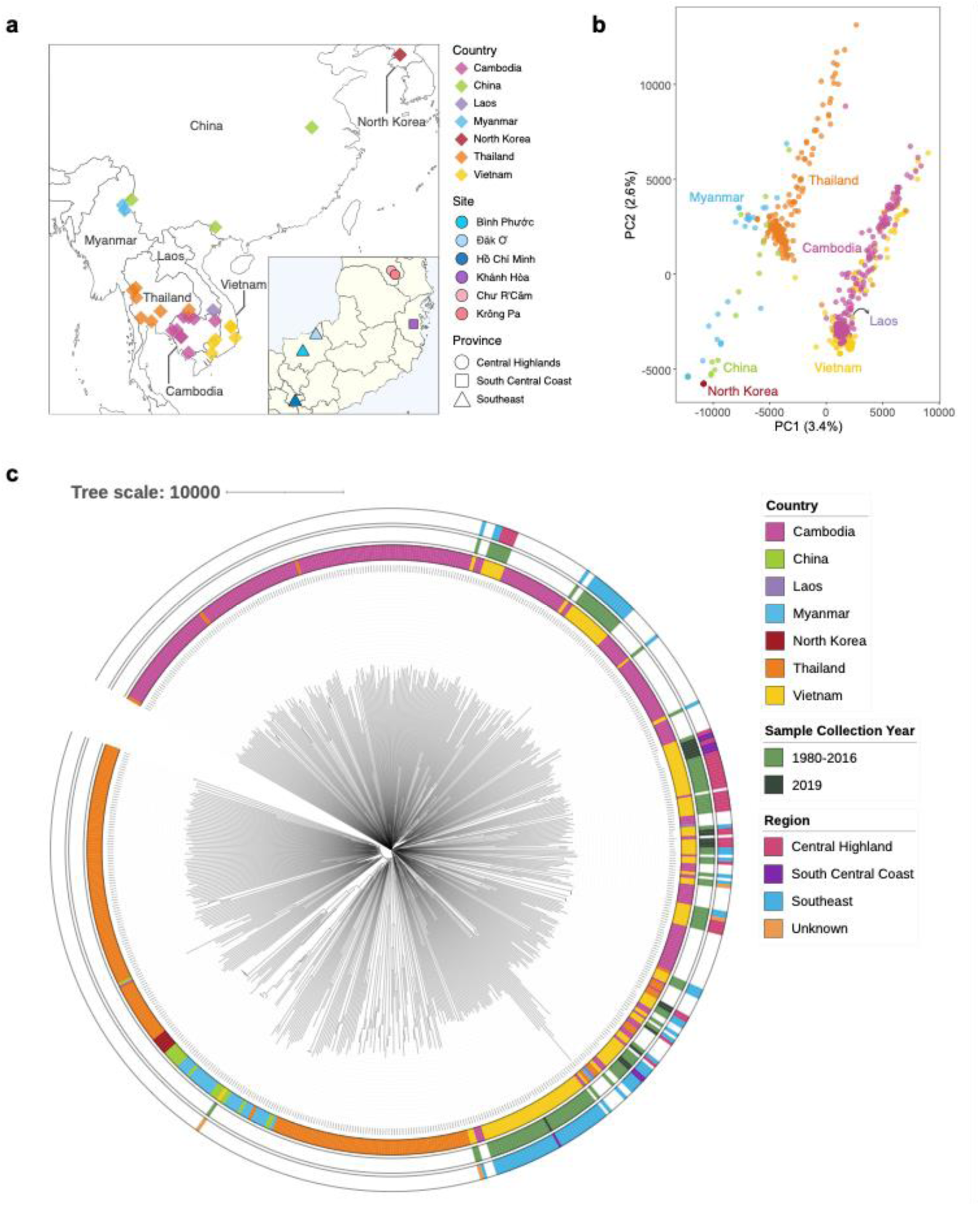
Population demographics and structure of 615 *P. vivax* isolates from around the GMS. **a)** Map showing the site of provenance of isolates, with those from within Vietnam displayed in an inset. **b)** PCA of the first two components showing two major geographical clusters of isolates. **c)** A neighbour-joining tree showing the relationship of isolates, revealing that Vietnamese isolates cluster by geography and not by year of collection.

Limited population structure was observed in Vietnamese isolates alone. In a SNP-based PCA coloured by province of origin (**Fig. S2a-b**), there appeared to be generalised clustering of isolates from Ho Chi Minh and Binh Phuoc, all from the South Central Coast, and loose clustering of isolates from Krông Pa. When coloured by year of collection (**Fig. S2c-d**), no distinct groupings were observed, suggesting geography is the main driver of the genetic landscape of Vietnamese *P. vivax*. These findings are in alignment with the SNP-based neighbour-joining tree (**Fig. 1c**), which showed random grouping of isolates in the outer strip coloured by year, but defined groupings in the outer strip coloured by region of provenance.

Across the GMS, *P. vivax* populations were structured in five distinct ancestral (*K*) pools, detected by ADMIXTURE analysis in populations with N≥10 (**Fig. 2**). An east-west axis was observed, where isolates from Cambodia and Vietnam shared ancestry through K1 and K5, and isolates from Myanmar and China shared ancestry through K2-K4. Isolates from Thailand appeared to be a bridge between the east and west, sharing ancestry with the eastern populations through K5, and with the western populations through K3 (**Table S3, Fig. S3**).

**Figure 2.**
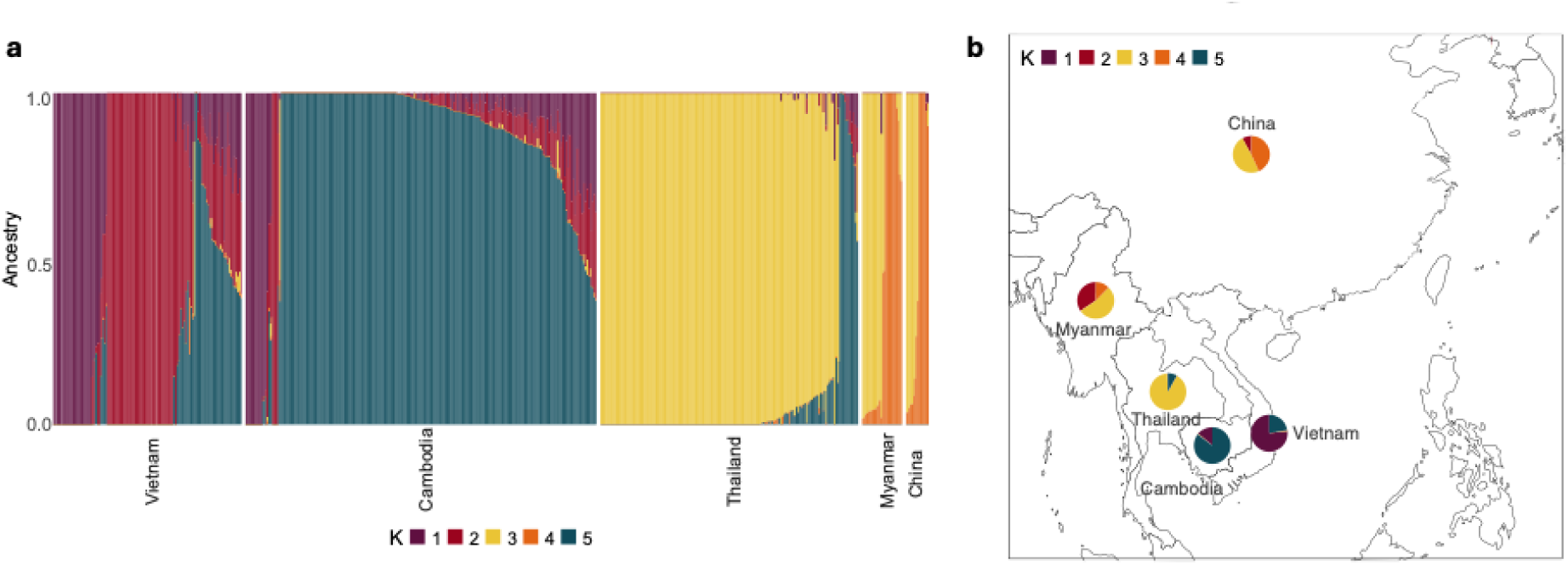
Population ancestry of 615 *P. vivax* isolates from around the GMS. **a)** Admixture estimation of *K = 5* ancestral populations from countries with ≥10 isolates, where each isolate is represented by a single bar coloured by proportion of each *K* population that makes up their ancestry. **b)** A map of the GMS showing, for each country, the distribution of the primary ancestral (*K*) population that isolates descends from.

#### Evidence of increasing relatedness in *P. vivax* populations in Vietnam

To infer *P. vivax* relatedness and gene flow in Vietnam, we estimated the proportion of genome fractions that shared identity by descent (IBD) among pairs of samples in the population using *hmmIBD*[31]. Relatedness is defined as the probability that, at any SNP, the two alleles drawn from the paired monoclonal *P. vivax* isolates are IBD, and therefore, have been inherited from a recently shared ancestor. Pairwise IBD values between 0.95-1 (95-100%) indicate recent shared ancestry. We retained sites with a minor allele frequency threshold of 0.01 (1%) or greater, and investigated monoclonal isolates exclusively, leaving 126,019 biallelic SNP sites from 86 samples for input into *hmmIBD*. The distribution of IBD values was highly skewed towards zero (**Fig. S4**) with the median proportion of pairwise IBD sharing at 0.0075 (0.75%). Of the 3,741 pairwise comparisons made, 16 had proportional IBD sharing >0.99, indicative of a clonal relationship, 5 had sharing indicative of full siblings, 18 had sharing indicative of half siblings, and 21 had sharing indicative of quarter siblings.

Exploration of the top 10% of sites sharing IBD within the 2009-2016 isolates revealed large tracts of DNA on chromosome 14 encompassing protein phosphatases 3 and 5 (1280kb-1300kb, IBD = 0.086), structural maintenance of chromosomes protein 2 (830kb-840kb, IBD = 0.102), nucleolar complex protein 4 (810kb-830kb, IBD = 0.128) (**Table S4**). Conversely, in 2019 isolates, chromosomes 1 and 13 exhibited excessive IBD proportions, including erythrocyte membrane-associated antigen (chr1:170kb-180kb, IBD = 0.136), protein phosphatase 12 (chr1:270-280kb, IBD = 0.134), protein phosphatase 11 (chr13:370kb-380kb, IBD = 0.146), and ABC transporter B family member 7 (chr13:410kb-420kb, IBD = 0146) (**Table S4**).

We next explored the wider spatial and temporal genetic structure of Vietnamese *P. vivax* by plotting IBD networks, where nodes represented a unique individual, and edges between nodes represented the degree of proportional IBD sharing at the level of quarter siblings, half siblings, full siblings, and clones. At the level of quarter siblings, 13 transmission clusters were observed, ranging from 2 to 10 members. Few spatially diverse clusters were observed, as we found that isolates from the same site or region of collection were often entangled in a network of connectivity (**Fig. 3a**). This was particularly evident in isolates from Gia Lai, where a network of 10 members was observed, and isolates from the Southeast region, with representation of 9 isolates from Binh Phuoc and Ho Chi Minh. Three inter-regional transmission clusters were observed between all regions; however, these clusters implicated one isolate from a different region with multiple isolates from the same region. For instance, a cluster of 8 individuals from the Southeast region was connected to one isolate from the South Central Coast. Investigating clonal transmission revealed 6 clusters with IBD ≥0.95, ranging between 2-4 members, and mainly involved isolates from the Central Highlands region (**Fig. 3a**). Inspection of temporal trends in IBD revealed that quarter siblings or closer (IBD ≥0.118) were derived from infections that occurred within the space of one to two transmission cycles (years), predominantly between 2015-2016. The longest identified transmission cluster was between 2014 and 2016. However, genealogical relationships of half-siblings to full clones revealed 3 persistent clonal lineages between 2015-2016, along with 3 clonal transmission clusters between isolates collected from 2019 (**Fig. 3b**). Overall, these findings indicate shared reservoirs of transmission at both a spatial and temporal scale.

**Figure 3.**
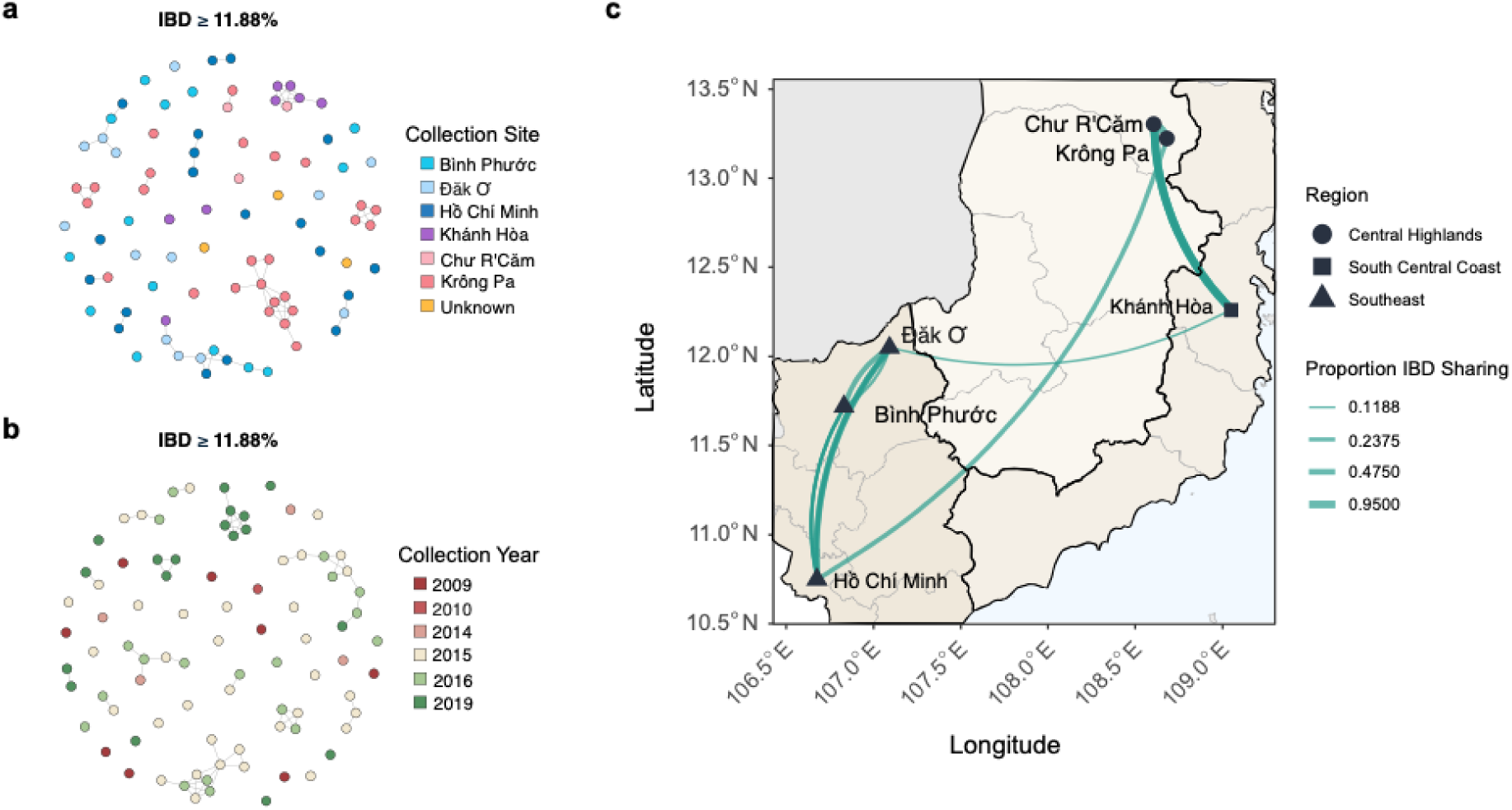
IBD networks inferring genetic relatedness of 86 monoclonal Vietnamese *P. vivax* isolates. Isolates are represented by nodes and coloured by **a)** site of collection or **b)** year of collection. Edges between nodes represent isolates that share IBD at a threshold equivalent to quarter siblings (IBD ≥ 0.1188). **c)** Proportional IBD sharing between isolates overlaid over geographic location to infer routes of parasite migration between the Central Highlands, South Central Coast, and Southeast regions of Vietnam. Here, nodes represent the geographic location of isolates within each region, and edges represent the proportion of IBD sharing equivalent to clones (IBD ≥ 0.95), full siblings (IBD ≥ 0.475), half siblings (IBD ≥ 0.2375) and quarter siblings (IBD ≥ 0.1188), with thickening edges symbolising increasing relatedness.

To further investigate parasite migration corridors and transmission routes, we mapped proportional IBD sharing across the Southeast, South Central Coast, and Central Highlands regions of Vietnam (**Fig. 3c**). Clonal transmission (IBD ≥0.95) was observed between Gia Lai and Khanh Hoa (Central Highlands–South Central Coast). We found our observation of clonal transmission and epidemiological connectivity between the Central Highlands and South Central Coast, potentially suggestive of a pathway for the spread of clonal lineages in Vietnam.

Similarly, parasites circulating in the Southeast region between Ho Chi Minh and Binh Phuoc were highly genetically related, with IBD proportions at the level of full siblings (IBD ≥0.475). These observations are in line with the principle of isolation by distance, which states that genetic similarity between populations increases as geographic distance between them decreases. Longer distance parasite migration was observed between isolates from Ho Chi Minh and Gia Lai, Binh Phuoc, and Khanh Hoa, however, these parasites were more genetically distinct, with IBD proportions at the level of full siblings and quarter siblings, respectively.

### Weak selection pressures on Vietnamese isolates from 2019

Genome-wide haplotype structure within monoclonal Vietnamese isolates was investigated to identify genomic loci under positive directional selection using the integrated haplotype score (*iHS*). The *iHS* identifies regions of high haplotype homozygosity relative to expectations under neutrality. Vietnamese isolates were explored in two distinct groupings: all publicly available isolates collected between 2009-2016 (N = 69), and newly sequenced isolates collected in 2019 (N = 18) (**Fig. 4a-b**). Across all Vietnamese isolates, hotspots of selection were found on chromosomes 3, 4, 6, 7, 10, and 14 (**Table S5**). These regions encompassed genes encoding proteins that are in direct interaction with the host immune response, including liver specific protein 2 (LISP2, PVP01_0304700), the merozoite surface proteins 4 and 5 (MSP4, PVP01_0418300; MSP5; PVP01_0418400), the Duffy binding protein (DBP, PVP01_0623800) and MSP1 (PVP01_0728900).

**Figure 4.**
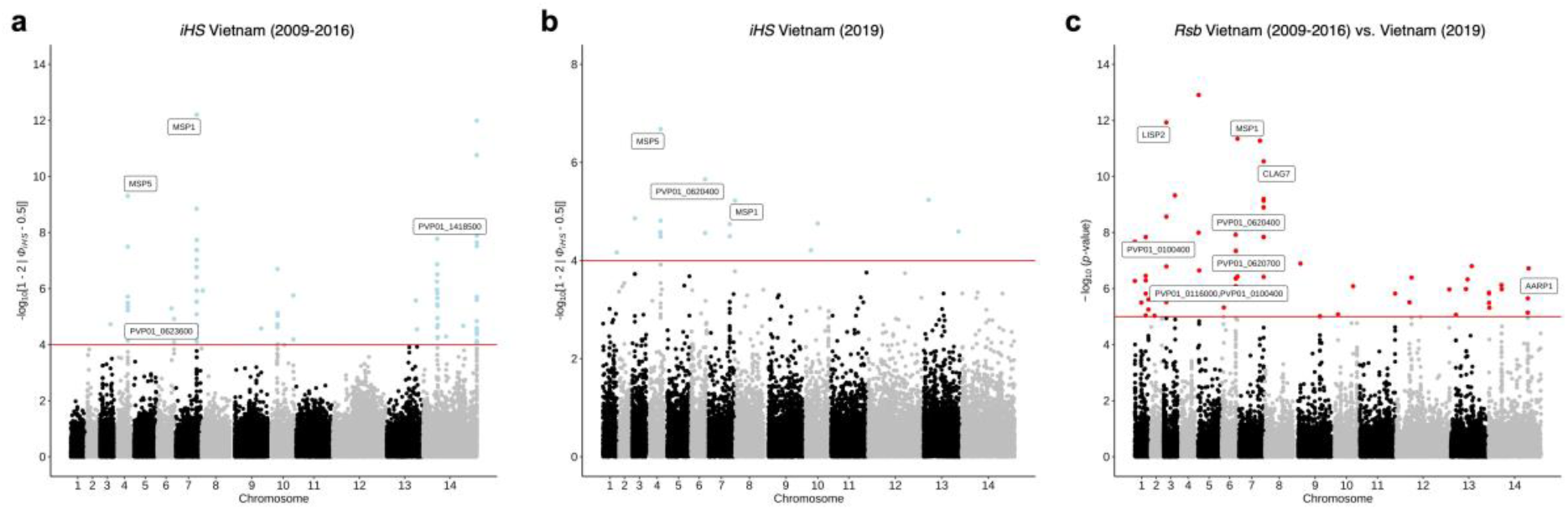
Evidence of recent directional selection in Vietnamese *P. vivax* isolates. The haplotype structure of Vietnamese isolates was analysed to identify regions of high local homozygosity relative to neutral expectation. Manhattan plots show integrated haplotype scores (*iHS*) for SNPs in the **a)** 2009-2016 population and **b)** 2019 population with critical loci defined as SNPs with an *iHS* score > 4.0. **c)** The cross-population statistic, *Rsb,* applied between the 2009-2016 vs. 2019 populations, showing hotspots of selection across the genome. Critical loci are defined with an *Rsb* value > 5.0.

Beyond these, other loci were gene products involved in replication and metabolism in the asexual stages, including splicing factors, assembly proteins, ribosomal proteins, and AP2 domain transcription factors. A region on chromosome 10 housing the multidrug resistance protein 1 (MDR1, PVP01_1010900), a putative mediator of chloroquine resistance, was also under directional positive selection. Many of the observations seen across the entire Vietnamese population were broadly consistent with findings in the 2009-2016 and 2019 populations. In the 2009-2016 population, hotspots of selection were observed on chromosomes 4, 6, 7, 10 and 14, with additional regions on chromosome 10, encompassing MSP3, and chromosome 14, encompassing *Plasmodium* helical interspersed sub-telomeric (PHIST) proteins. In the newly sequenced 2019 population, besides the MSPs on chromosomes 4 and 7, additional loci under directional positive selection were predominantly conserved *Plasmodium* proteins of unknown function.

The cross-population *Rsb* index was used to identify SNPs under selection in the 2019 population when compared with the earlier (2009-2016) population (**Fig. 4c**). The *Rsb* metric compares the extent of extended haplotype homozygosity between populations, with higher *Rsb* values ((-log10[p-value]) > 5.0) suggesting that a given allele is under positive selection in one population and not the other. A total of 168 SNPs were under differential selection between the 2009-2016 population when compared to the 2019 population (**Table S6**). Excluding the MSPs, SNPs were found in genes implicated in erythrocyte invasion, including cytoadherence linked asexual protein 7 (CLAG7, PVP01_0734500) and apical membrane antigen 1 (AMA1, PVP01_0934200). Other genes involved in life-cycle specific stages were also under differential selection, including the gametocyte development 1 gene (GD1; PVP01_0725700), which is critical for early sexual differentiation.

#### Temporal trajectories of allele frequencies in Vietnam between 2009-2019

To further investigate genetic differentiation between circulating Vietnamese *P. vivax* isolates over time, we used Hudson’s estimator of fixation index (F_ST_) for each SNP, specifically comparing 2009-2016 vs. 2019 populations. To reduce artefacts of population differentiation, we calculated Hudson’s F_ST_ on SNPs with a minor allele frequency of 1% and excluded sites where in either population, missing genotype calls were >20% (**Fig S5, Table S7**). When comparing the 2009-16 and 2019 populations, the genomic loci in the top 1% of F_ST_ values (>0.444, N = 24) were SNPs in genes encoding proteins that are involved in parasite development and survival such as GRP94 (PVP01_1440500, F_ST_ = 0.608), an endoplasmic protein stabilising proteins for export to host red blood cells[33], heat shock protein family members (PVP01_0108800, F_ST_ = 0.559), rhoptry associated proteins (PVP01_1144100, F_ST_ = 0.527), and DNA repair proteins (PVP01_1028500, F_ST_ = 0.450) (**Table 1**). Expanding our search to SNPs in the top 5% implicated the mutation G41S in AAT1 (PVP01_1120000, F_ST_ = 0.415), a chloroquine resistance mediator in *P. falciparum*[34]. Other loci in the top 5% F_ST_ values include genes involved in invasion or life-cycle specific stages, including MSP1 (PVP01_0728900, F_ST_ = 0.369), TRAP (PVP01_1218700, F_ST_ = 0.338), and LISP2 (PVP01_0304700, F_ST_ = 0.343), an early marker of liver stage development[35].

**Table 1.**
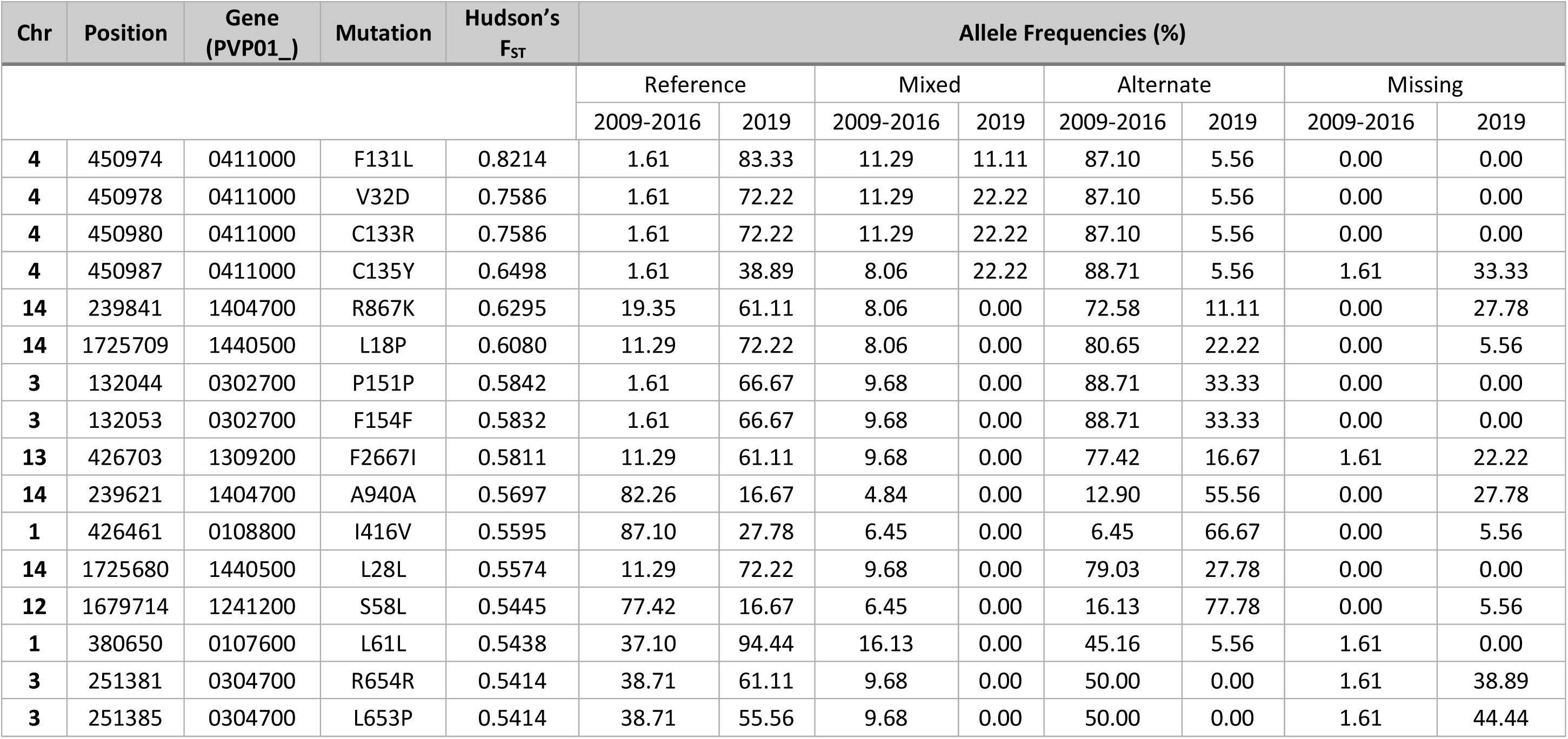
Allele frequencies of high F_ST_ SNPs in Vietnamese *P. vivax* 2009-2016 (N=115) vs. 2019 (N=18) populations.

| Chr | Position | Gene<br>(PVP01_) | Mutation | Hudson's<br>$F_{ST}$ | Allele Frequencies (%) | | | | | | | |
| --- | --- | --- | --- | --- | --- | --- | --- | --- | --- | --- | --- | --- |
|  |  |  |  |  | Reference |  | Mixed |  | Alternate |  | Missing |  |
|  |  |  |  |  | 2009-2016 | 2019 | 2009-2016 | 2019 | 2009-2016 | 2019 | 2009-2016 | 2019 |
| 4 | 450974 | 0411000 | F131L | 0.8214 | 1.61 | 83.33 | 11.29 | 11.11 | 87.10 | 5.56 | 0.00 | 0.00 |
| 4 | 450978 | 0411000 | V32D | 0.7586 | 1.61 | 72.22 | 11.29 | 22.22 | 87.10 | 5.56 | 0.00 | 0.00 |
| 4 | 450980 | 0411000 | C133R | 0.7586 | 1.61 | 72.22 | 11.29 | 22.22 | 87.10 | 5.56 | 0.00 | 0.00 |
| 4 | 450987 | 0411000 | C135Y | 0.6498 | 1.61 | 38.89 | 8.06 | 22.22 | 88.71 | 5.56 | 1.61 | 33.33 |
| 14 | 239841 | 1404700 | R867K | 0.6295 | 19.35 | 61.11 | 8.06 | 0.00 | 72.58 | 11.11 | 0.00 | 27.78 |
| 14 | 1725709 | 1440500 | L18P | 0.6080 | 11.29 | 72.22 | 8.06 | 0.00 | 80.65 | 22.22 | 0.00 | 5.56 |
| 3 | 132044 | 0302700 | P151P | 0.5842 | 1.61 | 66.67 | 9.68 | 0.00 | 88.71 | 33.33 | 0.00 | 0.00 |
| 3 | 132053 | 0302700 | F154F | 0.5832 | 1.61 | 66.67 | 9.68 | 0.00 | 88.71 | 33.33 | 0.00 | 0.00 |
| 13 | 426703 | 1309200 | F2667I | 0.5811 | 11.29 | 61.11 | 9.68 | 0.00 | 77.42 | 16.67 | 1.61 | 22.22 |
| 14 | 239621 | 1404700 | A940A | 0.5697 | 82.26 | 16.67 | 4.84 | 0.00 | 12.90 | 55.56 | 0.00 | 27.78 |
| 1 | 426461 | 0108800 | I416V | 0.5595 | 87.10 | 27.78 | 6.45 | 0.00 | 6.45 | 66.67 | 0.00 | 5.56 |
| 14 | 1725680 | 1440500 | L28L | 0.5574 | 11.29 | 72.22 | 9.68 | 0.00 | 79.03 | 27.78 | 0.00 | 0.00 |
| 12 | 1679714 | 1241200 | S58L | 0.5445 | 77.42 | 16.67 | 6.45 | 0.00 | 16.13 | 77.78 | 0.00 | 5.56 |
| 1 | 380650 | 0107600 | L61L | 0.5438 | 37.10 | 94.44 | 16.13 | 0.00 | 45.16 | 5.56 | 1.61 | 0.00 |
| 3 | 251381 | 0304700 | R654R | 0.5414 | 38.71 | 61.11 | 9.68 | 0.00 | 50.00 | 0.00 | 1.61 | 38.89 |
| 3 | 251385 | 0304700 | L653P | 0.5414 | 38.71 | 55.56 | 9.68 | 0.00 | 50.00 | 0.00 | 1.61 | 44.44 |

Except for *pvaat1*, no population differentiation effects were observed in any putative genes associated with antimalarial drug resistance, such as *pvmdr1, pvcrt-o,* or *pvmrp1*. To supplement this analysis, we investigated the prevalence of non-synonymous mutations in putative resistance loci between the 2009-2016 vs. 2019 populations (**Table 2**). By restricting our analysis to sites with <20% missingness within each population, this excluded all mutations in PvMRP1 (**Table S8**). Across PvMDR1, PvAAT1, PvK13, PvDMT1, and PvPPPK-DHPS, non-synonymous mutations were maintained at similar frequencies between the two temporal sub-populations. The greatest decrease in genotype frequency was in the G41S mutation in AAT1, a SNP with moderate population differentiation effects, going from 59.68% in 2009-2016 to 11.11% in 2019. The greatest increase in genotype frequency was the G196C mutation in PvPPPK-DHPS, a gene responsible for sulfadoxine resistance in the orthologue in *P. falciparum*, from 0% to 11.11% between 2009-2016 to 2019. No non-synonymous mutations were found in PvDHFR in either population. Overall, these observations suggest that allele frequency trajectories between 2009 and 2019 have not changed dramatically, and that broadly, the population is under similar selection pressures in 2019 as it was between 2009 and 2016.

**Table 2.** Allele frequencies of SNPs found within putatively associated antimalarial drug resistance genes (N=133)

| Gene<br>(PVP01_) | Gene name | Chr. | Position | Ref. | Alt. | Mut. | Allele Frequencies (%) |  |  |  |  |  |  |  |
| --- | --- | --- | --- | --- | --- | --- | --- | --- | --- | --- | --- | --- | --- | --- |
|  |  |  |  |  |  |  | Reference |  | Mixed |  | Alternate |  | Missing |  |
|  |  |  |  |  |  |  | 2009-2016 | 2019 | 2009-2016 | 2019 | 2009-2016 | 2019 | 2009-2016 | 2019 |
| 1010900 | MDR1 | 10 | 479908 | G | A | L1076F | 91.94 | 66.67 | 4.84 | 0.00 | 3.23 | 11.11 | 0.00 | 22.22 |
| 1010900 | MDR1 | 10 | 480601 | G | A | L845F | 70.97 | 50.00 | 4.84 | 0.00 | 24.19 | 27.78 | 0.00 | 22.22 |
| 1010900 | MDR1 | 10 | 481595 | A | T | S513R | 93.55 | 72.22 | 1.61 | 0.00 | 3.23 | 11.11 | 1.61 | 16.67 |
| 1120000 | AAT1 | 11 | 864055 | T | C | S25P | 83.87 | 94.44 | 4.84 | 0.00 | 9.68 | 5.56 | 1.61 | 0.00 |
| 1120000 | AAT1 | 11 | 864103 | G | A | G41S | 32.26 | 83.33 | 8.06 | 0.00 | 59.68 | 11.11 | 0.00 | 5.56 |
| 1120000 | AAT1 | 11 | 864355 | T | G | C125G | 1.61 | 11.11 | 1.61 | 0.00 | 95.16 | 83.33 | 1.61 | 5.56 |
| 1120000 | AAT1 | 11 | 864730 | G | A | E250K | 98.39 | 100 | 0.00 | 0.00 | 1.61 | 0.00 | 0.00 | 0.00 |
| 1120000 | AAT1 | 11 | 865571 | T | G | L530R | 98.39 | 100 | 0.00 | 0.00 | 1.61 | 0.00 | 0.00 | 0.00 |
| 1211100 | K13 | 12 | 485941 | G | A | S386N | 96.77 | 100 | 0.00 | 0.00 | 1.61 | 0.00 | 1.61 | 0.00 |
| 1340900 | PM4 | 13 | 1802108 | T | C | I165V | 24.19 | 16.67 | 8.06 | 0.00 | 67.74 | 61.11 | 0.00 | 22.22 |
| 1424900 | DMT1 | 14 | 1070933 | A | T | S310R | 9.68 | 11.11 | 1.61 | 0.00 | 88.71 | 88.89 | 0.00 | 0.00 |
| 1424900 | DMT1 | 14 | 1070943 | C | T | R307K | 96.77 | 100 | 0.00 | 0.00 | 3.23 | 0.00 | 0.00 | 0.00 |
| 1424900 | DMT1 | 14 | 1071124 | G | A | H247Y | 0.00 | 0.00 | 0.00 | 0.00 | 100 | 100 | 0.00 | 0.00 |
| 1424900 | DMT1 | 14 | 1071201 | A | G | I221T | 96.77 | 100 | 1.61 | 0.00 | 1.61 | 0.00 | 0.00 | 0.00 |
| 1424900 | DMT1 | 14 | 1071214 | C | T | V217I | 8.06 | 11.11 | 4.84 | 0.00 | 87.10 | 88.89 | 0.00 | 0.00 |
| 1424900 | DMT1 | 14 | 1071317 | G | C | C182W | 96.77 | 100 | 0.00 | 0.00 | 3.23 | 0.00 | 0.00 | 0.00 |
| 1424900 | DMT1 | 14 | 1071352 | T | G | K171Q | 98.39 | 100 | 0.00 | 0.00 | 1.61 | 0.00 | 0.00 | 0.00 |
| 1424900 | DMT1 | 14 | 1071643 | G | A | H74Y | 1.61 | 11.11 | 1.61 | 0.00 | 93.55 | 88.89 | 3.23 | 0.00 |
| 1424900 | DMT1 | 14 | 1071664 | G | T | L67I | 3.23 | 11.11 | 1.61 | 0.00 | 93.55 | 88.89 | 1.61 | 0.00 |
| 1429500 | PPPK-DHPS | 14 | 1270401 | G | C | A553G | 96.77 | 94.44 | 0.00 | 0.00 | 1.61 | 0.00 | 1.61 | 5.56 |
| 1429500 | PPPK-DHPS | 14 | 1270911 | C | G | G383A | 70.97 | 72.22 | 6.45 | 0.00 | 22.58 | 22.22 | 0.00 | 5.56 |
| 1429500 | PPPK-DHPS | 14 | 1271444 | C | T | M205I | 0.00 | 0.00 | 0.00 | 0.00 | 100.00 | 94.44 | 0.00 | 5.56 |
| 1429500 | PPPK-DHPS | 14 | 1271473 | C | A | G196C | 100 | 83.33 | 0.00 | 0.00 | 0.00 | 11.11 | 0.00 | 5.56 |

## Discussion

Elucidating parasite connectivity and transmission patterns using genomic data, particularly in countries and regions of low endemicity, is necessary for sustainable and maintained progress towards malaria elimination. Here, we performed an analysis of 18 newly sequenced and 115 publicly available whole-genome sequences from Vietnam to explore the evolving population and transmission dynamics of *P. vivax* between 2009-2019 as the country presses towards malaria elimination. Our analysis of the genetic trajectory of *P. vivax* within this period has uncovered focal inbreeding, weak population structure, and increasing relatedness in the population. Importantly, we found no profound change in allele frequencies in putative antimalarial drug resistance candidates between 2009-2019, suggesting Vietnam is primed for *P. vivax* elimination.

Despite their limitations, parasite-relatedness and multiplicity of infection are often a proxy indicator for transmission intensity[36,37]. In high transmission settings, *Plasmodium spp.* Infections with multiple distinct parasite clones are common as distinct parasite genotypes have more opportunities to recombine, whereas in low transmission settings, infections are often dominated by a single clone and are often highly related. We hypothesised that the isolates we collected in 2019 could either be entirely monoclonal or multiclonal infections of highly related clones (i.e., full siblings or closer). We found a pattern of decreasing within-sample diversity (F_WS_), with all 2019 isolates collected being monoclonal infections, evidencing the changing epidemiology of *P. vivax* in Vietnam, as intensified control interventions have led to a dramatic reduction in malaria transmission within this period. Several studies agree with this observation, including a study in West Papua, which showed that in a 14-year period where *P. vivax* incidence halved, the proportion of multiclonal infections also halved[38].

Genetic diversity of the Vietnamese *P. vivax* population was moderately high, consistent with a prior study in Central Vietnam[19], and in other low transmission settings, including Malaysia[39], Southwestern Pacific[40], and Panama[41]. This is likely due to the spatial heterogeneity of *P. vivax* transmission and is an indicator of highly focal inbreeding, where pockets of clonal lineages exist over long-distance spatial scales. This aligns with the epidemiology of *P. vivax* in Vietnam, which is heterogeneous and in decline, and marked by clusters of clonal infection hotspots, despite the overall malaria incidence in the country being very low.

Shifts in the evolutionary dynamics of the *P. vivax* parasite population can be observed using metrics such as IBD, which were compared here at a spatial and temporal scale to explore the impact of inbreeding in an era of population decline. Recent geostatistical modelling of residual malaria in Vietnam between 2019-2022 revealed considerable spatiotemporal heterogeneity in transmission, with highly receptive pockets in the Central Highlands and South Central Coast regions[42]. Based on our IBD analysis and clustering patterns in the genomic data, we found evidence of spatial aggregation of the Vietnamese *P. vivax* population, with clonal transmission especially evident in the Central Highlands and South Central Coast regions. Our findings are in alignment with prior studies of *P. vivax* populations in other low-transmission contexts, including Thailand[43], where some population structure was observed that separated isolates from western and eastern provinces, suggesting limited gene flow between parasite populations across the country. These findings agree with our infection multiplicity data, leading to the conclusion that recombination of Vietnamese *P. vivax* is occurring between very closely related individuals.

Evidence of the persistence of clonal lineages within Vietnam was limited to within the same malaria transmission season or from one season to the next. Our findings of clonal transmission of Vietnamese *P. vivax* are reminiscent of observations in prior studies of *P. vivax* in low transmission contexts, where a major or several minor clonal lineages persist over several transmission seasons, including in Malaysia[44] and Panama[41]. Along with our findings on geographic restriction of clonal lineages in Vietnam, our work again supports transmission of a moderately inbred *P. vivax* population. However, even across transmission seasons and within provincial boundaries, there are several index cases of isolates, including those collected in 2019, that are weakly related (quarter-siblings or less), adding a layer of complexity to the population structure.

We found little evidence of large-scale parasite migration in Vietnam. Most transmission connectivity we observed, inferred by IBD networks, was provincially confined, with 4 distinct clonal lineages circulating in the Central Highlands. There was only one instance of clonal transmission between the neighbouring provinces Central Highlands and South Central Coast. Although we have no data on the migration patterns of the human populations these isolates were derived from, the geographic patterns of relatedness we observed in the *P. vivax* population could, in part, be attributed to limited movement of people across different provinces due to geographical, socioeconomic, cultural, and/or work-related barriers. Taken together, these observations would support the highly monoclonal infections we observed in our IBD data, and provide some evidence of population stratification, where isolates from within the same region broadly clustered together. Importantly, it shows that there is limited gene flow between *P. vivax* sub-populations in Vietnam. Therefore, within transmission hotspots, recombination occurs between closely related individuals, if not clones. It is worth noting that the weakly related parasites observed here may represent residual pre-existing population variation or members of clonal lineages that persisted from earlier transmission cycles (e.g., 2017–2018) but were not sampled in this study.

In regions of low *P. vivax* endemicity, in addition to focal inbreeding, multiple circulating clonal populations or distinctive parasite genotypes can be a result of relapsing infections due to hypnozoite biology, or clonal expansion of imported cases from other regions, be it from within a country, from bordering countries, or beyond. It is well reported that the borders that Vietnam shares with Laos and Cambodia have a disproportionately higher incidence of malaria[5,45]. Economic migrants from malaria-endemic areas of Laos and Cambodia, along with mobile populations within the region, may further establish malaria in the country, leading to *P. vivax* parasites from Laos and Cambodia breeding with parasites in Vietnam. Our ADMIXTURE analysis of nuclear SNP data revealed shared ancestry of Vietnamese and Cambodian *P. vivax*, which has previously been described[11,12,15,46]. Although we observed no genomic evidence of cross-border transmission of malaria into Vietnam, we recognise that our sample sizes were inherently limited due to the rapidly declining case numbers of *P. vivax* in Vietnam and across Southeast Asia and highly heterogenous transmission cycles. Additionally, the weak relatedness between parasites we observed may also be from a population derived from asymptomatic individuals, which largely contributes to the infectious reservoir in low-endemicity settings, as seen in Ethiopia[47]. Many of the Vietnamese isolates in this study were sourced from provinces that are densely forested, hard to reach, and highly restricted, such as the Central Highlands. In these regions, *P. vivax* malaria disproportionately affects forest workers in labour-intensive industries (such as mining and plantations) and forest-going migrant populations[48]. Proximity to forests is a major risk factor for *P. vivax* malaria[45,49], as populations that reside or work outdoors, especially overnight, are more likely to have increased interactions with the mosquito vectors. Studies of the epidemiology of malaria in Central Vietnam frequently report a high proportion of asymptomatic *P. vivax* infections in forest-dwelling and forest-going individuals[5].

Chloroquine in conjunction with primaquine remains the front-line treatment for *P. vivax* in Vietnam. Although there is very limited evidence of chloroquine-resistant *vivax* in Vietnam, cases of treatment failure have been documented in the Binh Thuan[50], Quang Nam[51], and Gia Lai[52] provinces. While we observed no overall changes in the genotype frequencies between the 2009-16 vs. 2019 populations, we found the G41S mutation in AAT1, a mediator of chloroquine resistance in *P. falciparum*, with strong population differentiation effects, and a genotype frequency decreasing from 58.06% (2009-16) to 11.11% (2019). Although Hudson’s F_ST_ accounts for discrepancies in population sizes, given the limited sample size of the 2019 population, we can tentatively hypothesise that drug use during the higher transmission period of 2009-16 may have posed a significant selective pressure on AAT1. However, with intensified efforts that have successfully and drastically reduced *P. vivax* transmission, fewer infections, and thus, fewer treated infections mean fewer opportunities for drug-driven selection. In *P. falciparum*, the prevalence of resistant parasites is markedly lower in low transmission seasons, suggesting that drug-resistant parasites are less fit than drug-susceptible parasites[53,54]. Our findings here could point to this scenario in *P. vivax*, however, due to limited sample size and stochastic processes in low transmission settings, these findings may not be meaningful signals drug-driven selection and would therefore require further investigation, given that the presence of this SNP is private to countries in the GMS and Oceania[15].

To conclude, our findings provide a detailed analysis of the epidemiological and transmission context of *P. vivax* in three higher transmission regions in Vietnam between 2009-2019 as the country pushes towards elimination. Weak population structure, focal inbreeding, increasing infection monoclonality and relatedness, and limited temporal population differentiation define the *P. vivax* evolutionary trajectory within this 10-year period. Continued genomic surveillance alongside intensive control efforts will prove vital to ensure Vietnam meets its malaria elimination priorities.

## Data Availability

All raw Illumina sequence data used in this work is publicly available on the European Nucleotide Archive. Newly sequenced isolates from Vietnam are publicly available under Project PRJEB111293. Accession numbers and sample provenance for all samples are available in **Table S2.**

## Code Availability

All scripts used for population genetic analyses are publicly available at https://github.com/LSHTMPathogenSeqLab/malaria-hub.

## Acknowledgements

The study was funded by a joint Medical Research Council Newton UK – MOST Vietnam (MR/R026297/1) grant. G.C.N-J is funded by a BBSRC LIDo PhD studentship (BB/T008709/1). T.G.C and S.C are funded by UKRI MRC (IAA2129, MR/R026297/1, MR/X005895/1, and APP51212) and EPSRC (EP/Y018842/1) grants. The funders had no role in study design, data collection and analysis, decision to publish, or preparation of the manuscript.

## Author Contributions

N.T.H.B, S.C., N.Q.T., and T.G.C conceived and designed the study. N.T.H.B and N.T.H.N contributed samples. G.C.N-J conducted sample processing and DNA extraction. G.C.N-J and S.C. conducted sample sequencing. J.P provided software tools. G.C.N-J performed all bioinformatic analyses and interpreted results under the supervision of N.T.H.B, S.C., and T.G.C. G.C.N-J drafted the manuscript, with input from all authors. All authors reviewed and provided comments, approving of the final manuscript. G.C.N-J and T.G.C compiled the final submission.

